# The kinetic basis of CRISPR-Cas off-targeting rules

**DOI:** 10.1101/143602

**Authors:** Misha Klein, Behrouz Eslami-Mossallam, Dylan Gonzalez Arroyo, Martin Depken

## Abstract

Cas nucleases are popular tools for genome editing applications due to their ability to introduce DNA breaks at desired genomic locations. Such differential targeting is achieved through loading an RNA guide complimentary to the intended target sequence. As it turns out, sequences with only a partial match to the guide can also be cleaved. A large number of experiments have shed light on this off-targeting, outlining a number of rather peculiar empirical rules that detail the effect of mismatches at various positions and at various relative distances. We construct a kinetic model predicting on-target cleavage efficiency as well as off-target specificity. Our model explains a unified targeting rule for any target harboring mismatches, independent of their abundance and placing, and the observed decoupling between efficiency and specificity when protein-DNA interactions are weakened. We favorably compare our model to published experimental data from CRISPR-Cas9, CRISPR-Cpf1, CRISPR-Cascade, as well as to the human Argonaute 2 systems. Understanding the origins of off-targeting principles is important for the further development of CRISPR-Cas as a precise genome editing tool.

## INTRODUCTION

RNA guided nucleases (RGNs) target nucleic-acid sequences based on complementarity to any guide RNA (gRNA) loaded into the complex. This versatility, together with the ability to design synthetic gRNA complementary to any target of choice, holds great promise for gene editing and gene silencing applications (1–4). Among the known RGNs, the Clustered Regularly Interspaced Short Palindromic Repeats (CRISPR) associated (Cas) nucleases Cas9 (3, 5–8) and Cpf1 (Cas12a (9)) (10–12) are of special interest, as they are comparatively simple single-subunit enzymes. Cas nucleases have been successfully used to target sequences in genomes from a variety of organisms, ranging from prokaryotes to mammalian cells (7, 11–15). Recent success stories include the correction of point mutations relevant to human disease (16), the construction of sequence specific antimicrobials (17), the conveyance of infertility in malaria carrying mosquitoes (18), and the inactivation of retroviral elements to facilitate porcine-to-human organ transplants (19).

Cas nucleases originate from the CRISPR-Cas adaptive immune system, which many prokaryotes use to fight off foreign genetic elements. *In vivo,* the Cas protein (complex) is programmed by loading RNA transcribed from a CRISPR locus in the host genome. The transcribed sequence includes sections referred to as spacers, which were acquired during past encounters with foreign genetic elements (20–22). Once programmed, the Cas nuclease is able to target and degrade genetic elements with the same sequence as the stored spacer, and so offers protection against repeat invasions. Autoimmune response to sequences stored at the CRISPR locus is prevented through the additional requirement of a protein-mediated recognition of a short Protospacer Adjacent Motif (PAM) sequence present in the foreign genome, but not incorporated into the CRISPR locus with the spacer (23).

As viruses evolve in response to the selective pressure induced by the CRISPR immune system, the host is in turn under pressure to not only attack the target, but to also attack slightly mutated target sequences (24–26). It is therefore not surprising that Cas nucleases exhibit considerable off-target activity on sequences similar to the intended target (11, 12, 15, 27–34). Such off-target activity presents a severe problem for any therapeutic applications (4), as DNA breaks introduced at the wrong site could lead to loss-of-function mutations in a well-functioning gene, or the improper repair of a disease causing gene.

Surprisingly, the level of complementarity to the guide alone does not determine how strongly a particular sequence is targeted. To shed light on the determinants of off-target activity, a recent flurry of experiments has systematically probed the level of binding and/or cleavage on mutated target sequences (3), including high-throughput screens of large libraries of off-targets (11, 12, 14, 27, 29–31, 33–36), biochemical studies (6, 8, 10, 35, 37, 38), structural studies (39–45), and single-molecule biophysical studies (46–53) providing insights into the mechanics of targeting. To date, a number of rather peculiar targeting rules have been established for Cas nucleases: (i) *specificity-efficiency decoupling:* weakened protein-DNA interactions can improve target selectivity while still maintaining efficiency (54–57); (ii) *seed region:* single mismatches within a so-called seed region (a stretch of nucleotides following the PAM (58)) can completely disrupt interference, while mismatches further into the guide have much less of an effect (6, 8, 11, 12, 15, 27–34, 38, 47, 48, 50); (iii) *mismatch spread:* when mismatches are outside the seed region, off-targets with spread out mismatches are targeted most strongly (27, 30, 59). Although these experimental observations have already aided the development of strategies to improve the specificity of the CRISPR-Cas9 system (3, 54, 55, 57, 60, 61), an understanding of the mechanistic origin behind target selectivity is still lacking, and our ability to predict off-targets remains limited (1, 62).

Gaining a precise understanding of RGN specificity has the potential to greatly further therapeutic applications, as it could help both with the design of new enzyme re-engineering strategies for improved targeting and with choosing a gRNA that minimizes off-targeting (1, 62). Current computational approaches aimed at predicting off-targets for a given gRNA are often based on sequence alignment with the target, and discard potential targets if they have more than some (user-defined) threshold number of mismatches (62–65). To recover the mismatch-position dependence observed as seed regions (rule (ii)), such scoring schemes must be supplemented with phenomenological rules that penalize seed mismatches more than non-seed mismatches (66–68).

To move beyond *ad hoc* scoring schemes, we here use biophysical modelling to incorporate knowledge of the underlying targeting process. With this aim, it would be attractive to assume that the binding dynamics has had time to equilibrate before DNA degradation (69, 70), as this would allow us to use simple binding energetics to predict cleavage activity. Though attractive, this approach has recently been questioned by Bisaria *et al.* since the off-rate is generally not found to be much faster than the cleavage rate (71), as would be required for establishing a binding equilibrium before cleavage. In addition, the authors show how abandoning the equilibration assumption directly explains the specificity increase observed with shortened gRNA (60)—bypassing the need to fine tune energetic contributions along the guide.

Inspired by these observations, we go beyond binding energetics to build a biophysical model capturing the kinetics of guide-target hybrid formation. We show that also the targeting rules (i)-(iii) can be seen as simple consequences of kinetics. The targeting rules are captured by four parameters that pertain to transition barriers between metastable states of the RGN-guide-target complex, and we translate these into four experimentally observable quantities: the length of the seed region, the width of the transition region from seed to non-seed, the maximum amount of cleavage on single-mismatch off-targets, and the minimal distance between mismatches outside the seed region that allows for the cleavage of targets with multiple mismatches. By tying microscopic properties to biological and technological function we here to open the door to refined and rational reengineering of the CRISPR-Cas system to further its use in therapeutic applications.

Though we frame our considerations in terms of the well-studied and technologically important Cas9, our approach applies to any RGN that displays a progressive matching between guide and targeted (40, 47, 48, 50, 51) before cleavage (41, 72) (Figure 1A). To demonstrate the generality and power of our approach, we present fits to targeting data from Argonaute 2 (hAgo2), as well as type I, II and V CRISPR systems (11, 31, 38, 52).

**Figure 1:**
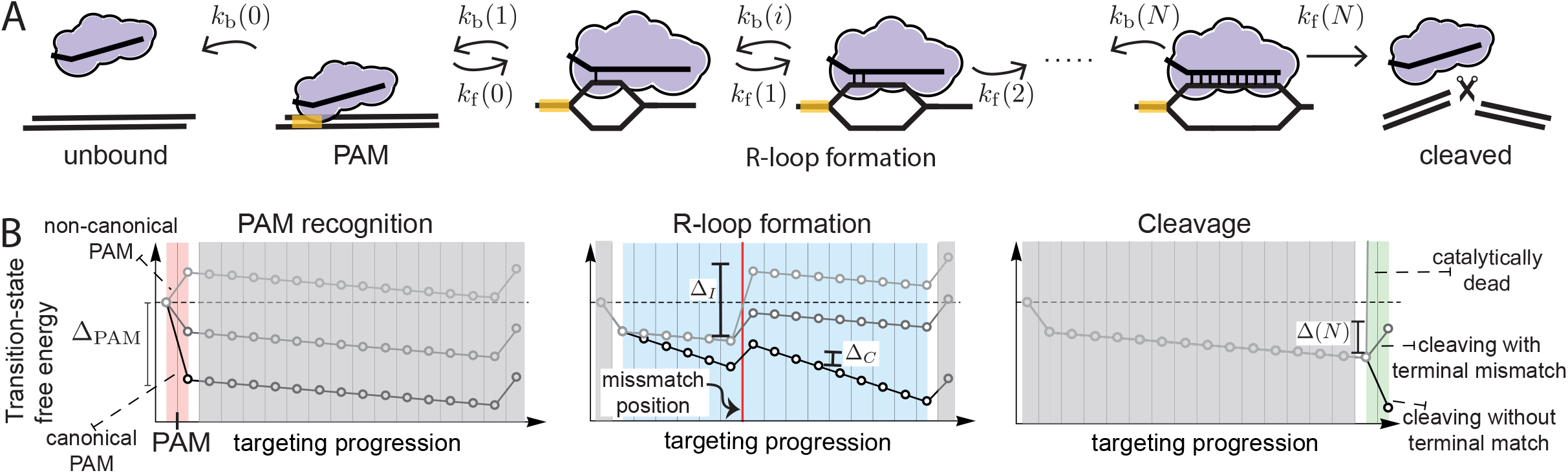
Kinetic model for target recognition by RNA guided nucleases. **(A)** The RGN binds its substrate at the PAM (state 0) from which it can unbind with backward rate *k_b_(0)* (solution state is −1) or form the first base pair of the R-loop (state 1) with forward rate *k_f_(0).* Next, the R-loop can grow (rate *k_f_(i)*) or shrink (rate *k_b_(i))* until the entire guide is bound (guide length *N*). Completion of the R-loop enables cleavage (rate *k_f_(N)*). In case of a non-PAM recognizing RGN, the solution state is given by state 0 and the RGN starts in state 1. **(B)** Our minimal model consists of, PAM recognition (red), who’s strength is set by *Δ_PAM_*. Matches within the R-loop (blue) bias its extension (*Δ_C_*), whereas any incorporated mismatches bias shrinking it (*Δ_I_*). The intrinsic catalytic rate (green) is determined through tuning the final (transition) state (*Δ_clv_*).

## RESULTS

At the start of target recognition, Cas nucleases bind to dsDNA from solution (Figure 1A). The subsequent recognition of a PAM sequence triggers the DNA duplex to open up, exposing the PAM proximal nucleotides to base pairing interactions with the guide. From here, an R-loop is formed, expanding a guide-target hybrid in the PAM distal direction (38, 40, 47–49, 51). If the target and guide reach (near) full pairing, cleavage of the two DNA strands is triggered (41, 72).

To establish the determinants of off-vs. on-target cleavage, we construct a biophysical model of sequential target recognition in the unsaturated binding regime (see **Methods**). Using this model, we can calculate the cleavage probability for any sequence and guide. To elucidate the mechanics of the targeting process, we envision it as a diffusion through a free-energy landscape, eventually leading to either unbinding from, or degradation of, the targeted sequence (**Figure S1**). The targeting decision can further be understood with reference to only a ‘transition landscape’ (Figure 1B), which show the transition states between any two metastable states, but excludes the free-energy of the (meta)stable states themselves (see Methods). In such a landscape, the R-loop typically grows whenever the forward barrier is lower than the backward barrier, or where the transition landscape tilts downward. To facilitate the discussion of our exact results, we use the rule-of-thumb (for justification see Methods) that a bound Cas9 is most likely to unbind before cleavage if the highest barrier to cleavage is greater than the largest barrier to unbinding, and vice versa (**Figure S1A-B**).

Though we treat the general scenario in the **methods** section, we here further limit ourselves to a minimal description with only four effective microscopic parameters, pertaining to the average kinetic bias for: R-loop initiation after PAM binding (Δ_PAM_), R-loop extension past a correctly matched (Δ_C_) and mismatched (Δ_I_) base pairs, and cleavage once the R-loop is fully formed (Δ_clv_) (for definitions see Figure 1B). Using this approach, we investigate to what extent our minimal model explains the three empirical targeting rules deduced from experiments.

### Rule (i): Specificity-efficiency decoupling

#### PAM recognition

Although PAM mismatches often completely abolish interactions with the target (27, 38, 50), binding to (and interference with) targets flanked by non-canonical PAM sequences has been observed (25, 26, 40, 46, 73, 74). Since PAM mismatches will shift the entire free-energy landscape upwards from the bound PAM state onwards (Figure 1B; left panel), these always increase the highest barrier to cleavage, thereby reducing the cleavage efficiency on any sequence. For increased specificity, we thus need the cleavage efficiency for the off-targets to be reduced more than for the target itself.

Protein reengineering approaches most easily affect the overall strength of PAM interactions, influencing the bias for both the correct PAM (Δ_PAM_) and incorrect 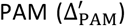. In Figure 2A we show the relative cleavage efficiency between incorrect and correct PAMs, and in Figure 2B we show the cleavage efficiency with the correct PAM—both as functions of the average kinetic bias 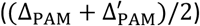 and the kinetic bias difference 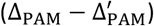. As long as the system operates in region A (Figure 2A), it is possible to increase the specificity by lowering the average kinetic bias toward R-loop formation, without changing the kinetic-bias difference. Outside this region, the system is either completely non-discriminating between PAMs (region C) or insensitive to the average kinetic bias (region B). Interestingly, it is only in region B that lowering the average bias also leads to a lower on-target efficiency (Figure 2B), and consequently the wild type (wt) nuclease can only be improved if brought into region A, where it is possible to engineer specificity increases without lowering the on-target efficiency. The transition-state diagrams shown in Figure 2C show a situation where the barrier to cleavage (right most node) is substantially lower than the barrier to unbinding (leftmost node) for two different PAM biases, resulting in near unit-probability cleavage for both, corresponding to region C in Figure 2A. Reengineering the nuclease to have an overall weaker PAM binding (Figure 2D) brings the system into region B, where the cleavage probability for the correct PAM (black) remains close to unity, while the probability of cleaving with the incorrect PAM (gray) is drastically lowered. The above scenario might explain how PAM mutant Cas9s are able to outperform their wildtype counterparts (55, 56) on specificity without losing efficiency.

**Figure 2:**
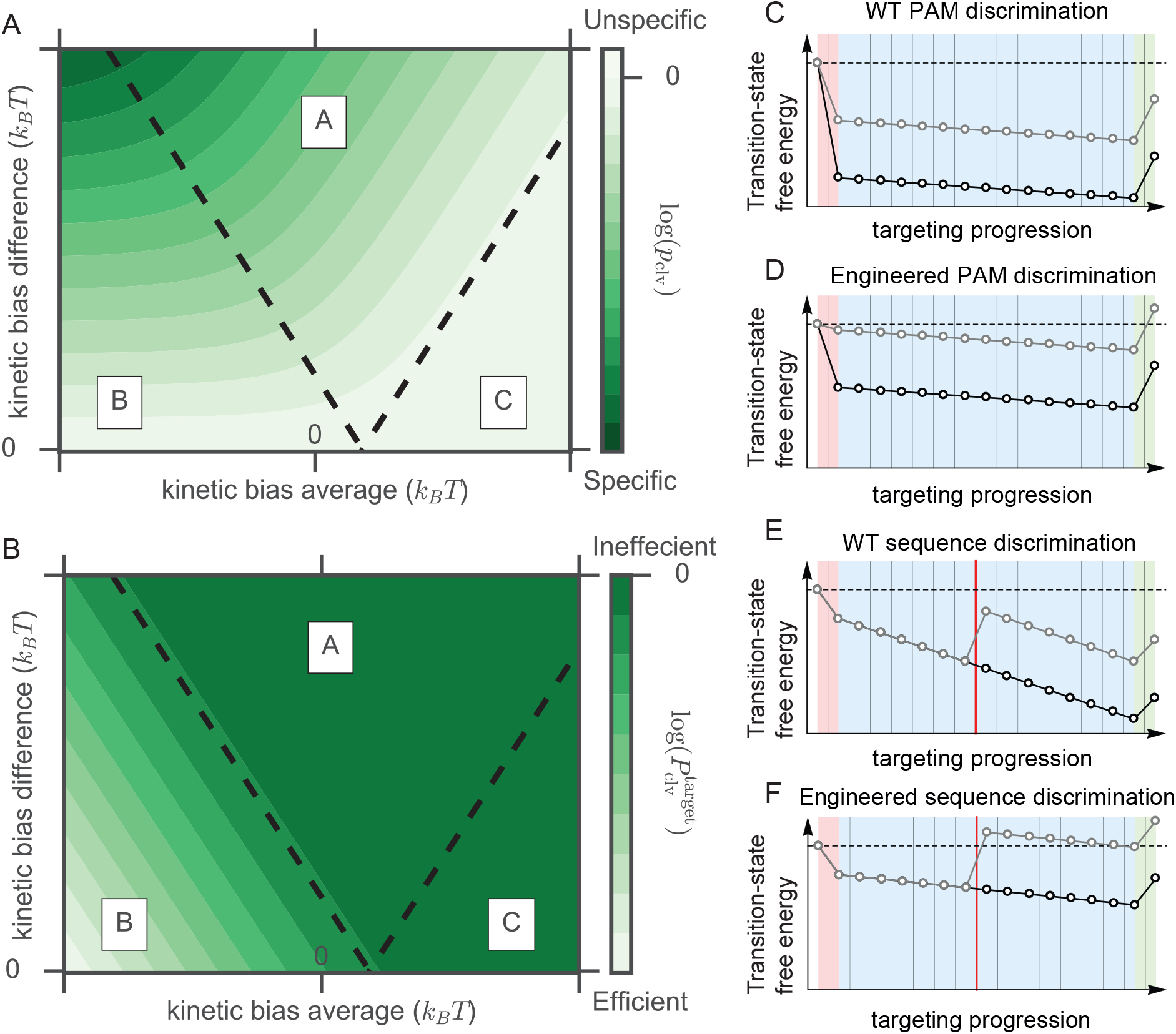
DNA-protein interactions affect target specificity. **(A)** Relative probability to cleave a target with a mismatched PAM, compared to one with a correct PAM, as a function of the average strength of PAM interactions and the difference in strength between the two PAMs. Independent of the sequence following both PAMs, taken to be identical, one can identify three regimes (S.I). In regime A, the RGN’s specificity is tunable through an overall decrease in PAM stability. (**B**) On-target efficiency for the target with the correct PAM. In regime A, the RGN’s efficiency is not compromised. **(C-D)** Weakening interactions between Cas protein and PAM allows the enzyme to reject **(D,grey)** previously cleaved **(C, grey)** mismatched targets, while maintaining a high on-target cleavage efficiency **(C-D, black)**. Similarly, weakening non-specific interactions **(E-F)** with DNA enhances the RGNs specificity **(E-F, grey)**, without rendering the enzyme to be inefficient **(E-F, black)**.

#### Sequence recognition

Another approach to gain specificity is to weaken the protein-DNA interactions effecting the bias for R-loop extension (54, 57). In **Figures 2E and F** we show how engineering the PAM-bound nuclease in this way, inducing a lower gain for correct base pairing, can render previously cleaved off-targets (gray line in Figure 2E) rejected (gray line in Figure 2F). In **Figures 2E and 2F** we further see how we can retain on-target specificity if the highest transition state towards cleavage (rightmost node of black line in Figure 2E and F) remains substantially lower than the transition state to unbinding (leftmost node of black line in Figure 2E and F). The above scenario might explain how mutant Cas9s could have an extended seed, while having negligible reduction in on-target cleavage activity (54, 57).

### Rule (ii): Seed region

#### PAM proximal mutations abolish cleavage

Following PAM binding, base pairing between guide and target is attempted (Figure 1B; middle panel). To establish if the experimentally observed dependence on the position of mismatches within the guide-target hybrid (6, 8, 11, 12, 15, 27–34, 38, 46–48, 50, 51) could originate from the kinetics of the targeting process, we calculate the relative cleavage probability on a sequence with a single mismatch positioned at *n*, compared to the cleavage probability on the target sequence. In the **Supplemental Information** we show that this relative cleavage probability is in general sigmoidal

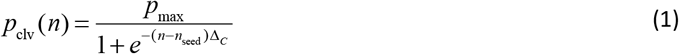

with *n*_seed_ the position where the cleavage probability is half that of its maximum *p*_max_ (Figure 3A). We identify *n*_seed_ as the length of the kinetic seed region, beyond which a mismatch will no longer strongly suppress cleavage (Figure 3A). From Equation 1 we see that the width of the transition from seed to non-seed region directly reports on the correct-match bias (Δ_C_, see Figure 1C, and **Methods**), becoming narrower as the bias increases (Figure 3A and **S2A**).

**Figure 3:**
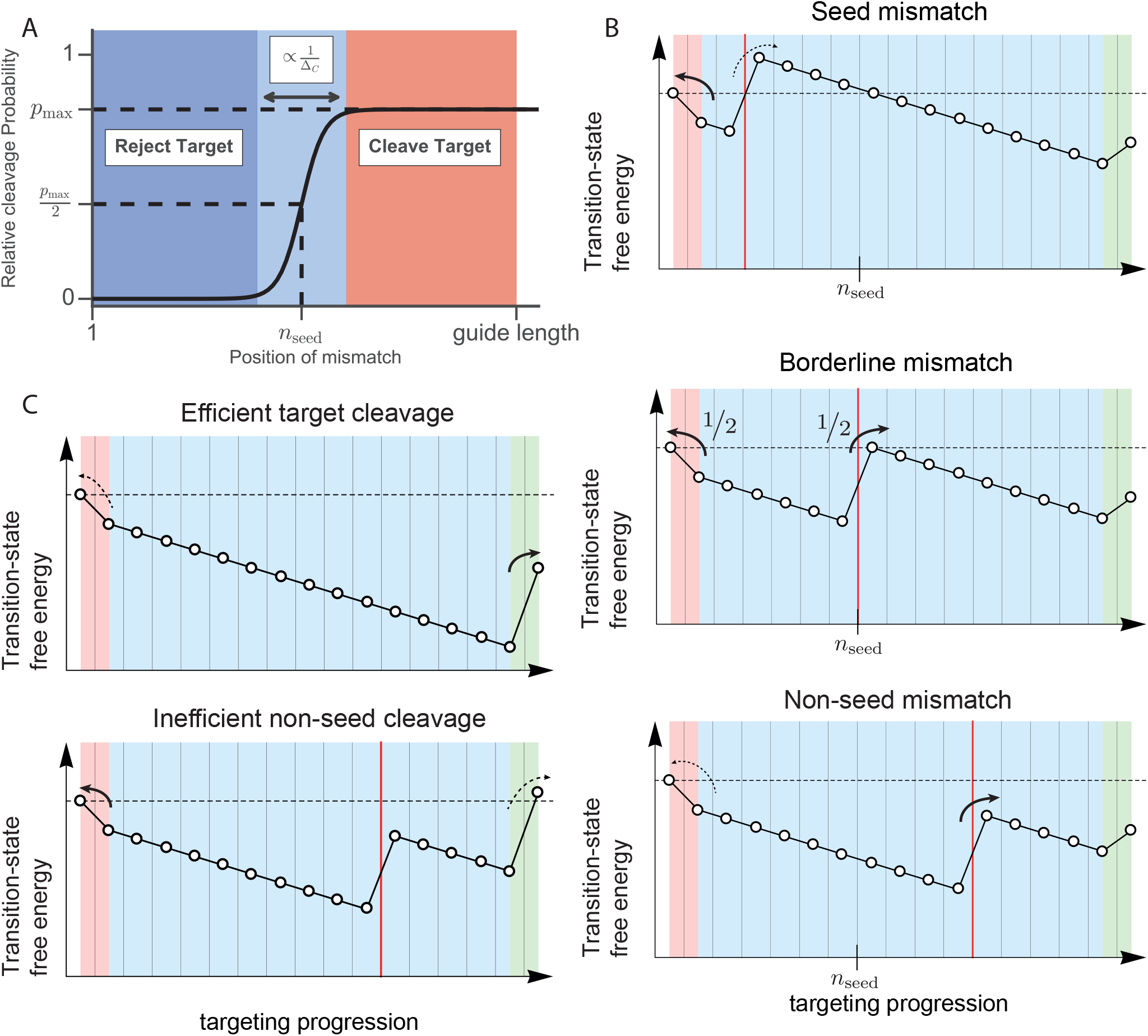
Single-mismatch off-targets. **(A)** Probability to cleave a target with a single mismatch compared to the completely complementary target. The microscopic parameters entering our minimal model (*Δ_C_*, *Δ_I_*, *Δ_PAM_*, *Δ_clv_*) result in a sigmoid with maximum off-target activity (*p_max_*), seed length (*n_seed_*) and width of the seed to non-seed transition (*Δ_C_*). **(B)** A functional seed length emerges as the site that, once mismatched, causes the effective barrier to overcome the mismatch to equal the one that hinders dissociation (*middle panel*). Placing the mismatch closer to the PAM favors dissociation (*top*), whereas the situation is reversed for PAM distal mismatches (*bottom*). **(C)** Slowing down the cleavage reaction (final straight line segment) can promote dissociation from targets with a PAM distal mismatch (*bottom*), while maintaining a high on-target activity (*top*).

The emergence of a seed-like region can be understood from considering the large-bias limit. When a mismatch is placed at *n*_seed_ (Figure 3B; middle panel), the highest barrier to cleavage matches the barrier towards unbinding, guaranteeing a near equal probability for cleavage and unbinding. Placing the mismatch closer to the PAM increases the highest barrier towards cleavage (Figure 3B; top panel), increasing the probability of rejecting such off-targets. Moving the mismatch distally from the PAM will gradually lower the highest barrier towards cleavage (Figure 3B; bottom panel), increasing the probability of accepting such off-targets. Though the exact form of the parameters of Equation 1 are given in the supplemental material, it is informative to also give the kinetic seed length in the large-bias limit (**Methods, S.I**),

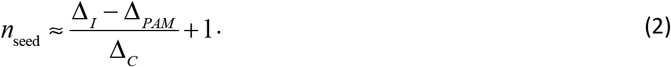

From this we see that PAM bias and the base pairing biases all contribute to setting the extent of the seed region (Figure 3A, **S2B**). Weakening the PAM or correct-match bias extends the seed region, while weakening the bias for incorrect matches shrinks it.

After PAM recognition and R-loop formation are completed, cleavage completes the targeting process (Figure 1B; right panel). Tuning the final transition state allows us to toggle between different regimes of minimal single-mutation specificity. Targets with a PAM distal mismatch get cleaved with near unity probability (*p*_max_ ≈ 1) if all transition states towards cleavage (including the cleavage step) lie well below the transition state to unbinding (Figure 3B; upper panel, **S.I**). For slow enough enzymatic activity, the final barrier towards cleavage never goes far below the barrier to unbinding (Figure 3C; lower panel), limiting the maximal cleavage compared to the perfect match (*p*_max_ < 1) (Figure 3C; lower panel). Consequently, there can be a noticeable effect on off-target activity also when the mismatch is outside the seed region (Figure 3A, **S2C**). Reversing this logic implies that a *p*_max_ < 1 is indicative of a relatively slow cleavage reaction.

#### Differential binding versus differential cleavage

Catalytically dead systems (for example dCas9 or Cascade without Cas3) bind strongly to sites that their catalytically active counterparts do not cleave (14, 28, 33, 34, 36, 59). In order to explain this effect, we model inactive systems with a very large cleavage barrier (gray in Figure 1B; right panel). **Figures S3A and S3B** show the dissociation constant (**Methods**) for targets harboring a single mismatch. In agreement with experimental observations (38), our model predicts a dissociation constant that is higher when a mismatch is placed closer to the PAM.

Similar to the cleavage efficiency in the kinetic regime, the equilibrium dissociation constant takes on a sigmoidal form (**S.I**). However, the resulting seed length (**Figure S3A**) is different from its kinetic counterpart resulting from Equation 2 (**S.I.**). Binding affinities therefore do not need to report on differential cleavage activities. In general, the gene editing (Cas9) and gene silencing (dCas9) capabilities should be seen as two separate properties of the RGN. For example, the most stable configuration of the RGN on the mismatched target shown in Figure 4A is a bound state with a partial R-loop (purple). However, a catalytic active variant will most likely reject this off-target (gray) as the barrier to cleavage is higher than to unbinding. Hence, even though cleavage sites are strong binders (Figure 4A; right panel), observing a long binding time on an off-target site should not be taken to imply that this site will also display substantial off-target cleavage (Figure 4A; left panel).

**Figure 4:**
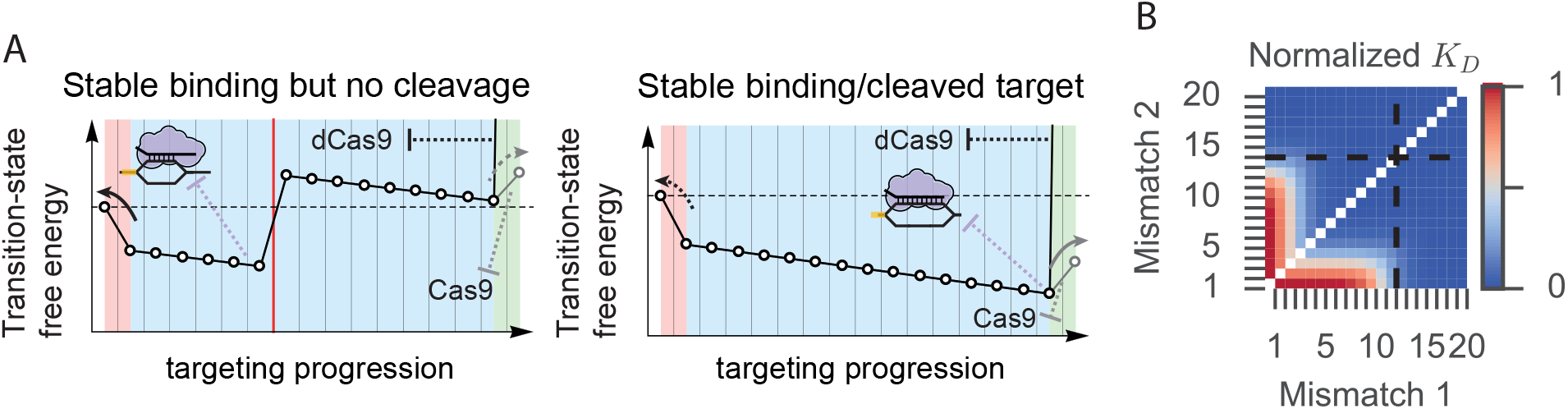
Differential binding versus differential cleavage. **(A)** When bound to the cognate target (*right*), a catalytically dead RGN (*black*) is most likely stably bound and has formed the complete R-loop (*purple*). It’s catalytically active counterpart will likely cleave this target (*grey*). However, on mismatched targets (*left),* a dCas9 protein (*black)* can be stably bound after forming a partial R-loop (*purple)* even though the catalytically active variant would likely reject this target (*grey*). **(B)** Dissociation constant for targets with any combination of two mismatches. Seed length indicated with dashed lines. (*δ_PAM_* =7.5k_B_ T, *δ_l_* =8k_B_ T,*δ_c_* =1k_B_ T).

### Rule (iii): Mismatch spread

#### Spreading mismatches over non-seed region promotes cleavage

Next, we consider more complex mismatch patterns, starting by addressing all possible dinucleotide mismatches (Figure 4B and 5A). The overall patterns obtained strongly resemble experimental observations (27, 30, 59). As expected, placing both mismatches within the seed disrupts cleavage (Figure 5A). However, moving both outside the seed does not necessarily restore cleavage activity. With the first mismatch outside the seed region, a second mismatch only abolishes cleavage if it is situated before *n*_seed_ + *n*_pair_ (Figure 5B). In the large-bias limit (**Methods, S.I**):

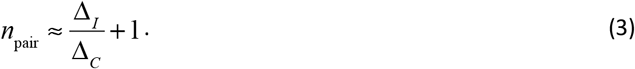

**Figure 5:**
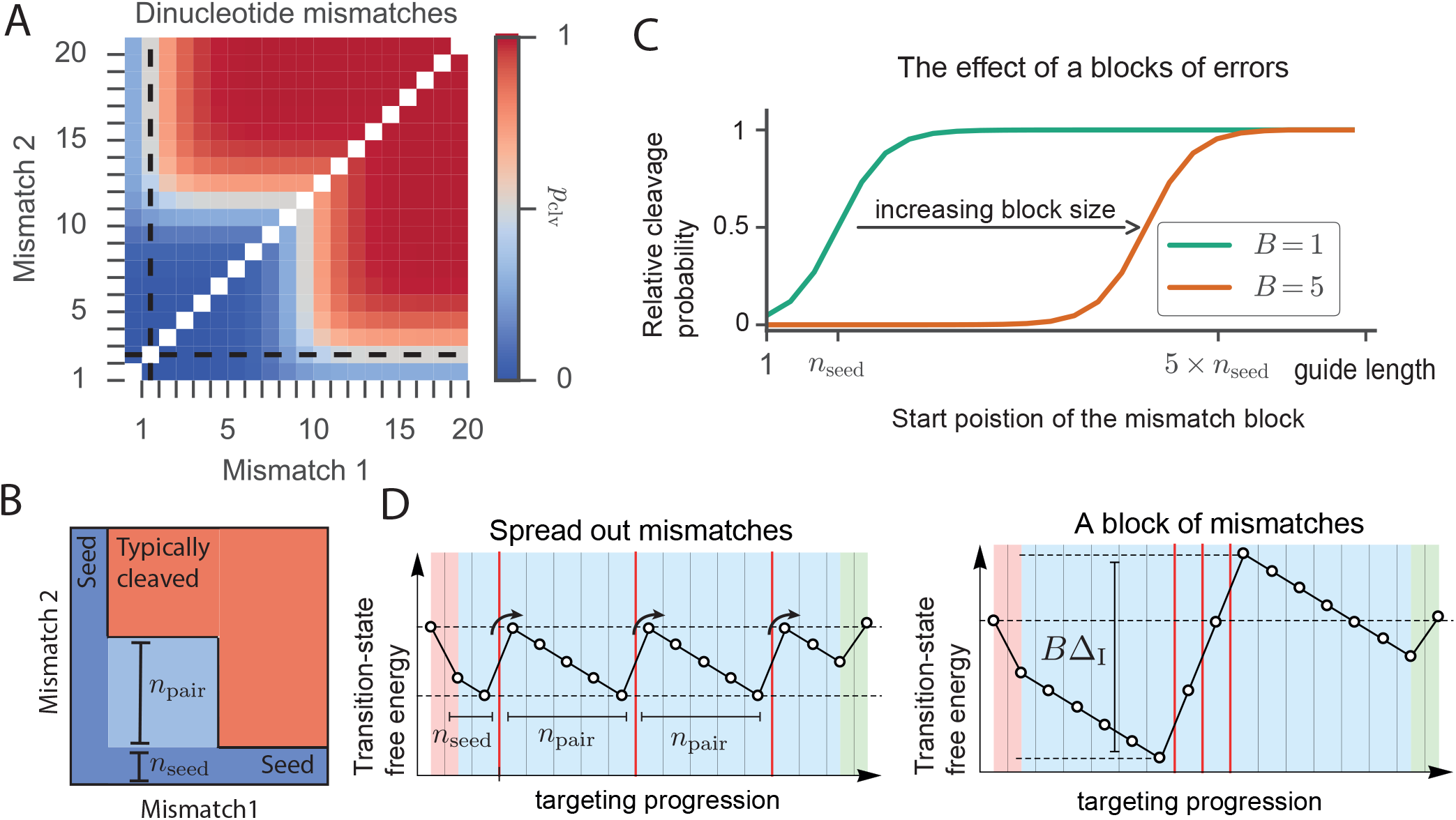
Multiple mismatches. **(A)** Relative probability to cleave a target with two mismatches. Seed length (*n_seed_*) indicated with dashed lines. (*Δ_PAM_*=1k_B_T, *Δ_l_* =4k_B_T,*Δ_C_*=1k_B_T, *Δ_clv_*=−100k_B_T). **(B)** Schematic view of probability to cleave a target with two mismatches. Placing both mismatches within the seed abolishes interference (*dark blue*). Placing both PAM distal restores interference (*red*). Placing one mismatch outside the seed restores interference if the second mismatch is placed beyond the indicated light blue area. Hence, if the first mismatch is placed at the seed’s border, a minimal distance of *n_pair_* between mismatches is needed. **(C)** Probability to cleave a target with a stretch (‘block’) of mismatches (size *B*) as a function of the location of the first mismatch (*Δ_PAM_*=3.5k_B_T, *Δ_l_* =4k_B_T,*Δ_C_*=1k_B_T, *Δ_clv_*=-100k_B_T). **(D)** Spreading the mismatches out (placing them *n_pair_* apart) (*left*) favors cleavage compared to a stretch of mismatches (*right*).

The general form of the two-mismatch seed region is shown in Figure 5B, where only off-targets in the red region lead to cleavage. In the dark blue region, off-targets are rejected due to the first mismatch, and in the light blue region they are rejected due to the second mismatch.

The single- and double-mismatch rules can now be unified and generalized (see Figure 5D; left panel) into a single rule for any number of mismatches:

> *Off-targets will typically be rejected if any mismatch, say the m:th mismatch, is positioned closer than n*_seed_ + (*m* – 1)*n_pair_ from the PAM*.

Note that for systems without PAM, *n*_seed_ equals *n*_pair_ in the above generalized targeting rule. The above rule also captures the extreme case of a ‘block’ of *B* consecutive mismatches, which has also been investigated experimentally (49, 52). Placing such a block effectively acts as placing a single mismatch with the bias Δ_I_ scaled by the size of the block (Figures 5C, 5D and **S4**), giving a block seed region of size *n*_seed_ + (*B* – 1)*n*_pair_.

### Comparison to experimental data for a broad class of RNA guided nucleases

To test our model, we acquired published datasets acquired for different RGN systems, and fitted Equation 1 to singly mismatched targets and blocks of mismatches. The fitted sigmoid has only three effective fit parameters (*p*_max_ or *K*_D,max_, *n*_seed_, and Δ_C_), so we can unfortunately not get an estimate for all microscopic parameters from the single-mismatch datasets (**S.I**)—for this, further experiments are required, as outlined below. Details of the fitting procedure and additional fits can be found in the **Supplemental Information**.

Perhaps the best characterized RGN system is the Type-II CRISPR associated *Streptococcus Pyogenes* Cas9 (spCas9). The dataset from Anderson *et al.* (31) traces out the sigmoidal trend particularly well. For this data set we fit out a kinetic seed of about 11 nt (68% conf. interval [11.0,11.4]), and an average bias per correct base pair of about Δ_C_ = 1.7 *k*_B_*T* [1.15,4.0)) (Figure 6A, **S5**). This bias is about a factor two lower than the average gain per correct base pair for RNA:DNA hybrids in solution (75, 76), indicating that association with the RGN destabilizes the hybrid. This is in line with recent studies demonstrating that the protein has a strong contribution to the energetics of the resulting bound complex (53, 54, 57). The relative cleavage probability levels-off around *p*_max_ = 0.74 [0.72,0.77], indicating that spCas9 retains some specificity even against errors that are outside the seed.

**Figure 6:**
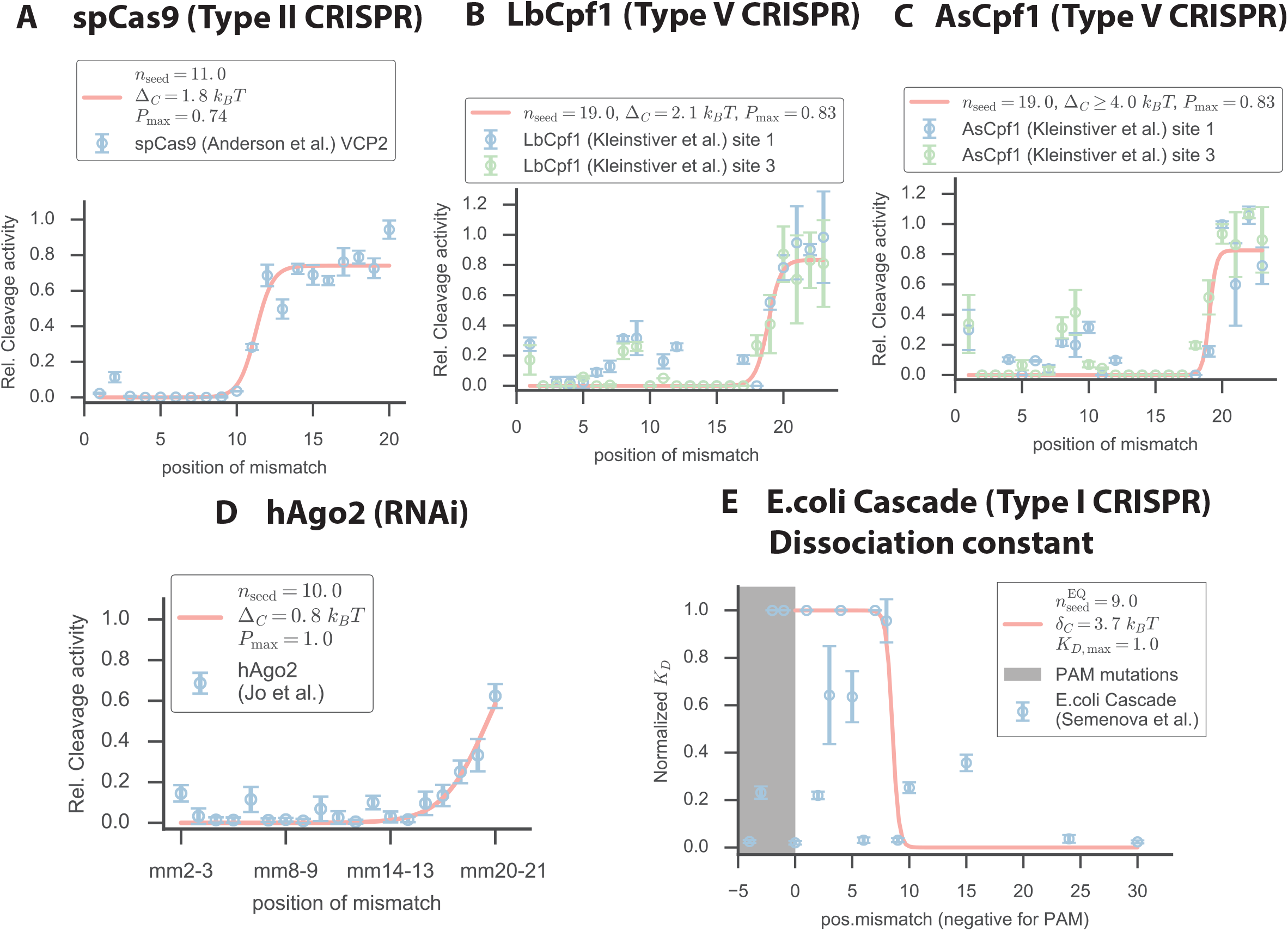
Comparison to experimental data. Fit of sigmoid (Eqn 1) to experimental data from (A) spCas9 (Anderson et al., ref 31) (B) LbCpf1 (Kleinstiver et al., ref 11) (C) AsCpf1 (Kleinstiver et al., ref 11) (D) Human Argonaute 2 (Jo et al., ref 52) (E) E.coli Cascade complex (Semenova et al., ref 38) Reported values corresepond to the median of 1000 bootstrap replicates (**S.I, Figure S5-S8**). Reported intervals in the text correspond to 68% confidence intervals.

Recently, the type V CRISPR associated enzyme Cpf1 has been characterized as another single-subunit RGN (10). Kleinstiver *et al.* (11) performed *in vivo* (human cells) cleavage assays using two different variants named LbCpf1 (Figure 6B,**S6**) and AsCpf1 (Figure 6C,**S7**). Both variants exhibited quantitatively similar off-targeting, both with seed lengths (*n*_seed_ ≈ 19 nt,LbCpf1:[18.5,19.2], AsCpf1:[18.7,19.3]) and maximum off-target activity (*p*_max_ ≈ 0.8, LbCpf1:[0.66,1.0], AsCpf1:[0.71,1.0]). Although, the stability of the RNA:DNA hybrid within the enzymes (as measured by Δ_C_) differ in the two cases, which mainly affects their respective on-target efficiencies, the difference is not statistically significant given the data at hand (LbCpf1:[1.15,4.0), AsCpf1:[2.25,4..0)). Compared to spCas9, the Cpf1s are much more specific as the seed region is significantly larger.

Single-molecule FRET experiments done with hAgo2 (52) utilized targets with two consecutive mismatches. Given that hybrid formation is not preceded by a PAM-like interaction, and that consecutive mismatches impose a combined penalty (Figure 5C and D), the estimated half-saturation point is approximately twice the kinetic seed length for a single mismatch (*n*_seed_ ≈ 10 nt [9.5,9.9]). The hAgo2 data thus suggests a similar seed length as that of spCas9 (Figure 6D,**S8**), consistent with the observation that hAgo2 and spCas9 display structural similarities within their respective seed regions (41). Our fits further reveal that hAgo2 likely exhibits a substantially lower gain per correctly formed base pair (Δ_C_≈ 0.7 *k*_B_*T* [0.63,0.92]).

Unlike the aforementioned RGNs, the Type I CRISPR uses a multi-subunit protein complex, termed Cascade, to target invaders (77). Semenova *et al.* (38) measured the dissociation constant *in vitro* of the *E.Coli* (subtype IE). Fitting their data, we find that mismatches within the first 9 nt of the guide lead to rapid rejection (Figure 6E). Interestingly, the energetic gain for a match again suggests a large contribution of the protein to the overall stability, similar to the other CRISPR systems tested (Δ_C_ ≈ 3.7 *k*_B_*T*).

## DISCUSSION AND CONCLUSION

We have presented a general description of target recognition by RNA guided nucleases with a progressive matching between guide and target (Figure 1A)—describing both CRISPR and Argonaute systems. In its simplest form, our model contains only two parameters to describe the R-loop formation process: an average bias towards incorporation beyond a match (Δ_C_) and an average bias against extending the R-loop beyond a mismatch (Δ_I_) (Figure 1B; middle panel). Despite the simplifications going into this minimal model, we can quantitatively understand the targeting rules for these RGNs as resulting from kinetics (specificity-efficiency decoupling, Figure 2C–F; seed region, Figure 3B; and mismatch spread, Figure 5D). Moreover, our model also explains why there is in general a poor match between cleavage propensity and binding propensity for these nucleases (14, 28, 33, 34, 36, 59) (Figure 4 D). Based on our model we have been able to establish a general targeting rule: *Off-targets will typically be rejected if any mismatch, say the m:th mismatch, is positioned closer than n*_seed_ + (*m* – 1)*n*_pair_ *from the PAM* (see Equation 2 and 3).

Although Figure 6 shows that our model can already describe experimental data from various RGNs, the number of microscopic parameters in the physical model (Δ_PAM_, Δ_C_, Δ_I_, and Δ_clv_, see **Methods** and Figure 1B) exceeds the number of fit parameters available from single-mismatch experiments (Δ_C_, *p*_max_, and *n*_seed_) (**S.I**). It is therefore not possible to determine all the microscopic parameters from single-mismatch experiments alone. However, **Figures 5B** shows that with two mismatches, we could also fit out n_pair_, and so determine all the microscopic parameters. Alternatively, we show in the **S.I.** that a combination of data from single-mismatch binding and cleavage experiments can provide all four microscopic parameters as well. It should be possible to directly extract all four microscopic parameters once such extended datasets become available.

Interestingly, the data from Cpf1 (Figure 6B and C) shows an increased tolerance to mismatches of nucleotides 1,2,8 and 9 compared to our minimal model, and a second independent study shows the same behavior (12). A recent structural study of LbCpf1 mentions nucleotide 9 being solvent exposed, suggesting a lack of involvement in the interference process (39). However, the available crystal structure of AsCpf1 does not suggest any similar feature (44). Additional work is presently underway to fully understand such non-monotonically increasing cleavage probabilities from a physical-modelling perspective.

The great diversity of systems showing the same basic behavior, hints towards and overarching physical principles governing RNA/DNA guided RNA/DNA targeting. Still, CRISPR type I-E systems have been observed to exhibit a second non-canonical binding mode onto targets with mutated PAM and or seed sequences (46, 78), which enables rapid introduction of immunity to mutated sequences (24–26). Additional research is needed in order to couple the observed PAM/seed dependent and independent binding modes. Also, Type-III CRISPR systems do not use PAM recognition to discriminate self from non-self. Instead, complementarity with the direct repeats flanking both sides of the spacer sequence in the host CRISPR locus is used (79). Perhaps, such systems do not bind their targets in a unidirectional fashion. Our general modeling approach could still be applicable, and it should be possible to tie dynamics to energetics and structure also for these systems.

It will further be interesting to see if newly discovered CRISPR and Argonaute systems (81, 82) use the same or different targeting principles to the systems described here. Intriguingly, a novel bacterial Argonaute system (82) seems to be tolerant to mutations of the first few nucleotides, despite having part of its guide nucleotides arranged in a similar fashion to the seed region of a known Ago variant.

The current formulation only deals with the problem of recognizing a target once it is at hand. Under physiological conditions, the target sequence must first be located amongst the large pool of available binding sites within the cell. This target search process constitutes a long standing problem in biological physics and the precise mechanism of ensuring both speed and specificity is still under debate (80). Another effect not taken into account thus far is the concentration of active RGNs, which could be an important further extension as *in vitro* work is often done at high concentrations compared with *in vivo* studies (27, 32).

In conclusion, our kinetic model is capable of explaining the observed off-target rules of CRISPR and Argonaute systems in simple kinetic terms. After having established the general utility of this approach, the next step will be to move beyond our minimal model and allow for a dependence on both the nature of matches/mismatches and their positions. Fitting such a generalized model against training data could improve on present target prediction algorithms, as it uses a minimal set of parameters to capture the basic targeting rules deduced from experiments.

## METHODS

### A general model for RGNs with progressive R-loop formation followed by cleavage

Given the observed dependence of cleavage activity on Cas9 concentration (15, 28, 32, 33), we here limit ourselves to the regime where nuclease concentrations are low enough that all binding sites are unsaturated. The unsaturated regime is also the regime with the highest specificity, and should therefore be of particular interest in gene-editing applications.

We define the cleavage efficiency P_clv_(*s*|*g*) as the fraction of binding events to sequence *s* that result in cleavage, given the RGN is loaded with guide sequence *g*. If we in the unsaturated regime assume the binding rate to be independent of sequence, we can express the relative rate of non-target vs. target cleavage as

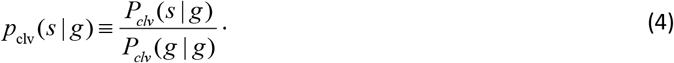

This relative efficiency is a direct measure of *specificity*, approaching unity for non-specific targeting (*P*_clv_(*s*|*g*) ≈ *P*_clv_(*g*|*g*)) and zero for specific targeting (*P*_clv_(*s*|*g*) ≪ *P*_clv_(*g*|*g*)).

In our model, we denote the PAM bound state as 0 and the subsequent R-loop states by the number of base pairs that are formed in the hybrid, 1,..., *N*. Each of the states *n* = 1,..., *N* are taken to transition to state *n* – 1/*n* + 1 with backward/forward hopping rate *k*_b_(*n*)/*k*_f_(*n*) (Figure 1A). The ratio between forward and backward rates sets the relative probability of going forward and backward from any state, and can be parametrized in terms of Δ(*n*), the difference in the free-energy barrier between going backwards and forwards from state *n* (see **Figure S1A, S.I**),

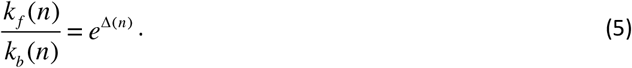

Here we measure energy in units of *k*_B_*T* for notational convenience, and we will refer to Δ(*n*) as the bias toward cleavage. The model (Figure 1A) is known as a birth-death process (83, 84), and the cleavage efficiency is given by the expression (**S.I**),

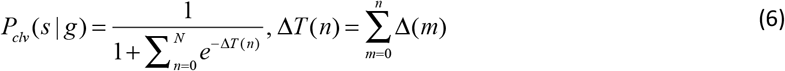

Here Δ*T*(*n*) represents the free-energy difference between the transition-state to solution (**Figure S1A**) and the forward transition state from position *n* (**Figure S1A-B**).

For systems like hAgo2, there is no initial PAM binding (80, 85, 86), and the sums in Equation 6 should omit the PAM state (*n*, *m* = 0). It is therefore convenient to separate out the bias due to PAM binding (Δ_PAM_ = Δ(0)), and describe both CRISPR and non-CRISPR systems explicitly

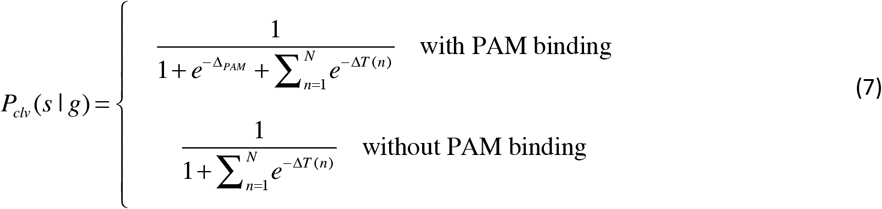

### Building intuition by using the transition landscape (large bias limit)

As the cleavage probability is completely determined by the set of relative transition-state free-energies, we can simplify the free-energy diagrams by omitting the metastable states and draw ‘transition landscapes’ by connecting the transition states by straight lines. A downward slope (Δ> 0) of such a connection means a bias to move forward (extend hybrid), while an upward slope (Δ< 0) means a bias to move backward (shrink the hybrid) (see Equation 5 and **Figure S1A**).

Though we will use the exact results of Equation 7 for all calculations, it is useful to build intuition for the system by considering the case of large biases. In this limit, the term (say *n* = *n*\*) with the highest transition-state dominates the sum in Equations 6 and 7 (**Figure S1A-B**), and the cleavage efficiency can be approximated as

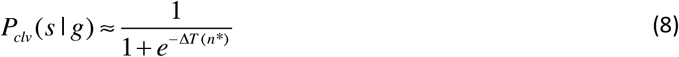

Based on this we deduce the rule-of-thumb that cleavage dominates (*P*_clv_ > 1/2) if the first state of the transition landscape is the highest (Δ*T*(*n*\*) > 0) (**Figure S1A**). Conversely, a potential target is likely rejected (*P*_clv_ < 1/2) if any of the other transition states lies above the first (Δ*T*(*n*\*) < 0) (**Figure S1B**).

### A minimal model for RGNs with progressive R-loop formation followed by cleavage

Given that the defining feature of RGNs is their programmability to target any sequence, we expect the major targeting mechanisms to depend more strongly on mismatch position than on the precise nature of the mismatches. With this in mind, we consider a sequence independent model with the aim of finding a description that captures the gross, sequence averaged, features with a minimal number of parameters.

Focusing first on how PAM binding effects the system (Figure 1B; left panel), we see that Δ(0) = Δ_PAM_ controls the bias between initiating R-loop formation and unbinding. A canonical PAM (black) promotes R-loop initiation, while a non-canonical PAM lessens (darker gray) or reverses (lighter gray) the bias towards R-loop formation. Note that PAM independent systems omit this initial step.

Turning to the bias of R-loop progression, we represent the guide-target hybrid as a sequence of matches (*C*, correct base pairing) and mismatches (*I*, incorrect base pairing). Defining the average bias *towards/against* extending the R-loop by one *correct/incorrect* base pair as Δ_C_/Δ_I_ (Figure 1B; middle panel), we take Δ(*n*) = Δ_C_ or Δ(*n*) = –Δ_I_ depending on if the base pairing is correct or incorrect (**S.I**). In the middle panel of Figure 1B we show a transition landscape with moderate gains for correct base pairings and moderate costs for incorrect base pairings (dark gray). The black transition landscape corresponds to an increased gain for matches, while the light gray corresponds to an increased penalty for mismatches. Cleavage activity has also been observed on off-targets with small insertions and deletions (37). We can easily capture the effect of such indels by assuming that they induce bulges in the guide-target hybrid. Bulges come at an energetic cost, and can therefore be included by allowing variable mismatch biases Δ_I_.

Lastly, considering the bias between cleavage and unwinding of the R-loop, we assume that an incorrect base-pair at the terminal position adds the same change in bias as it did in the interior of the R-loop. Therefore, introducing the cleavage bias Δ_clv_, we take Δ(*N*) = Δ_C_ – Δ_clv_ and Δ(*N*) = –Δ_I_ – Δ_clv_ for the terminal base being correct or incorrect respectively (Figure 1B; right panel). In the right panel of Figure 1B, we show an example of the terminal bias Δ(*N*) with a terminal match (black), terminal mismatch (dark gray), and for a catalytically dead nuclease (light gray).

### Dissociation constant for catalytically dead nucleases

Apart from examining cleavage propensity, many experiments have focused on the binding of catalytically dead Cas9 (dCas9) or other catalytically dead RGNs (28, 33–35, 38, 51, 59) (Figure 1B; right panel, light gray line). To be able to relate pure binding experiments to cleavage experiments, we also calculate the dissociation constant *K*_D_ for our minimal model when describing a catalytically dead system (Δ_clv_≈ –) (**S.I., Figure S1C**) through

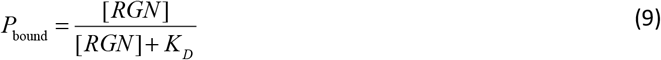

Here *P*_bound_, equals the probability to bind a substrate in any of the (*N*) possible R-loop configurations and follows from Equation 7 (**S.I**). Further, [RGN] denotes the concentration of effector complex. In the **Supplemental Information** we show that we can describe the binding of dCas9 with three parameters, out of which two are identical to R-loop biases (Δ_C_ and as Δ_I_) for the catalytically active system.

## ACKNOWLEGDEMENTS

First and foremost, MD would like to thank Ralph Seidel for introducing him to the problem of sequential target recognition during a visit to Delft. We would also like to thank Aafke van den Berg, Michiel Bongaerts and Kristian Blom for fruitful discussion regarding the theoretical modeling. We further acknowledge the great discussions we have had with all members of the CRISPR/microRNA community in Delft: Viktorija Globytė, Luuk Loeff, Tao Ju (Thijs) Cui, Stanley Chandradoss, Sungchul Kim, Seung Hwan Lee, Chirlmin Joo, Rebecca McKenzie, Jochem Vink, Sebastian Kieper and Stan Brouns. MK also thanks Iason Katechis for introducing him to the Python fitting library (lmfit).

Rebecca, Thijs and Chirlmin for their comments on the manuscript.

We gratefully thank Emily Anderson, Benjamin Kleinstiver and Keith Joung, Sungchul Hohng, Soochul Shin and Myung Hyun Jo as well as Ekaterina Semenova for sharing their data and answering all the questions we had.

This work was supported by the Netherlands Organization for Scientific Research (NWO/OCW), as part of the Frontiers in Nanoscience program. MD acknowledges financial support from a TU Delft startup grant.

## AUTHOR CONTRIBUTIONS

M.K and M.D designed the research. M.K, D.G.A and B.E.M performed the research and, together with M.D, interpreted the data. M.K and M.D wrote the paper, with input from the other authors.

## References

1. Tycko J, Myer VE, Hsu PD (2016) Methods for Optimizing CRISPR-Cas9 Genome Editing Specificity. Mol Cell 63(3):355–370.

2. Wu X, Kriz AJ, Sharp PA (2014) Target specificity of the CRISPR-Cas9 system. Quant Biol 2(2):59–70.

3. Hu JH, Davis KM, Liu DR (2016) Chemical Biology Approaches to Genome Editing: Understanding, Controlling, and Delivering Programmable Nucleases. Cell Chem Biol 23(1):57–73.

4. Cox DBT, Platt RJ, Zhang F (2015) Therapeutic genome editing: prospects and challenges. Nat Med 21(2):121–131.

5. Gasiunas G, Barrangou R, Horvath P, Siksnys V (2012) Cas9-crRNA ribonucleoprotein complex mediates specific DNA cleavage for adaptive immunity in bacteria. Proc Natl Acad Sci U S A 109(39):E2579–86.

6. Jinek M, et al. (2012) A programmable dual-RNA-guided DNA endonuclease in adaptive bacterial immunity. Science (80-) 337(6096):816–821.

7. Mali P, et al. (2013) RNA-guided human genome engineering via Cas9. Science (80-) 339(6121):823–826.

8. Cong L, et al. (2013) Multiplex Genome Engineering Using CRISPR/Cas System. Science (80-) 339(February):819–824.

9. Makarova KS, Zhang F, Koonin E V. (2017) SnapShot: Class 2 CRISPR-Cas Systems. Cell 168(1–2):328–328.e1.

10. Zetsche B, et al. (2015) Cpf1 Is a Single RNA-Guided Endonuclease of a Class 2 CRISPR-Cas System. Cell 163(3):759–771.

11. Kleinstiver BP, et al. (2016) Genome-wide specificity profiles of CRISPR-Cas Cpf1 nucleases in human cells. Nat Biotechnol (June):1–7.

12. Kim D, et al. (2016) Genome-wide target specificities of Cpf1 nucleases in human cells. Nat Biotechnol 34(8):876–881.

13. Cho SW, Kim S, Kim JM, Kim J-S (2013) Targeted genome engineering in human cells with the Cas9 RNA-guided endonuclease. Nat Biotechnol 31(3):230–2.

14. Duan J, et al. (2014) Genome-wide identification of CRISPR/Cas9 off-targets in human genome. Cell Res 24(8):1009–1012.

15. Fu Y, et al. (2013) High-frequency off-target mutagenesis induced by CRISPR-Cas nucleases in human cells. Nat Biotechnol 31(9):822–826.

16. Komor AC, Kim YB, Packer MS, Zuris JA, Liu DR (2016) Programmable editing of a target base in genomic DNA without double-stranded DNA cleavage. Nature 61(16):5985–91.

17. Bikard D, et al. (2014) Exploiting CRISPR-Cas nucleases to produce sequence-specific antimicrobials. Nat Biotechnol 32(11):1146–50.

18. Hammond A, et al. (2016) A CRISPR-Cas9 gene drive system targeting female reproduction in the malaria mosquito vector Anopheles gambiae. Nat Biotechnol 34(1):78–83.

19. Yang L, et al. (2015) Genome-wide inactivation of porcine endogenous retroviruses (PERVs). Science (80-) 350(6264):1101–4.

20. Wiedenheft B, Sternberg SH, Doudna J a. (2012) RNA-guided genetic silencing systems in bacteria and archaea. Nature 482(7385):331–338.

21. Sorek R, Lawrence CM, Wiedenheft B (2013) CRISPR-mediated adaptive immune systems in bacteria and archaea. Annu Rev Biochem 82(May):237–266.

22. Hsu PD, Lander ES, Zhang F (2014) Development and applications of CRISPR-Cas9 for genome engineering. Cell 157(6):1262–1278.

23. Westra ER, et al. (2013) Type I-E CRISPR-Cas Systems Discriminate Target from NonTarget DNA through Base Pairing-Independent PAM Recognition. PLoS Genet 9(9). doi:10.1371/journal.pgen.1003742.

24. Datsenko K a., et al. (2012) Molecular memory of prior infections activates the CRISPR/Cas adaptive bacterial immunity system. Nat Commun 3(May):945.

25. Fineran PC, et al. (2014) Degenerate target sites mediate rapid primed CRISPR adaptation. Proc Natl Acad Sci U S A 111(16):E1629–38.

26. Künne T, et al. (2016) Cas3-Derived Target DNA Degradation Fragments Fuel Primed CRISPR Adaptation Correspondence Cas3-Derived Target DNA Degradation Fragments Fuel Primed CRISPR Adaptation. Mol Cell 63:1–13.

27. Hsu PD, et al. (2013) DNA targeting specificity of RNA-guided Cas9 nucleases. Nat Biotechnol 31(9):827–32.

28. O’Geen H, Henry IM, Bhakta MS, Meckler JF, Segal DJ (2015) A genome-wide analysis of Cas9 binding specificity using ChIP-seq and targeted sequence capture. Nucleic Acids Res 43(6):3389–3404.

29. Fu BXH, St. Onge RP, Fire AZ, Smith JD (2016) Distinct patterns of Cas9 mismatch tolerance *in vitro* and *in vivo*. Nucleic Acids Res:gkw417.

30. Fu BXH, Hansen LL, Artiles KL, Nonet ML, Fire AZ (2014) Landscape of target: Guide homology effects on Cas9-mediated cleavage. Nucleic Acids Res 42(22):13778–13787.

31. Anderson EM, et al. (2015) Systematic analysis of CRISPR-Cas9 mismatch tolerance reveals low levels of off-target activity. J Biotechnol 211:56–65.

32. Pattanayak V, et al. (2013) High-throughput profiling of off-target DNA cleavage reveals RNA-programmed Cas9 nuclease specificity. Nat Biotechnol 31(9):839–43.

33. Kuscu C, Arslan S, Singh R, Thorpe J, Adli M (2014) Genome-wide analysis reveals characteristics of off-target sites bound by the Cas9 endonuclease. Nat Biotechnol 32(7):677–683.

34. Wu X, et al. (2014) Genome-wide binding of the CRISPR endonuclease Cas9 in mammalian cells. Nat Biotechnol 32(7):670–676.

35. Ran FA, et al. (2015) In vivo genome editing using Staphylococcus aureus Cas9. Nature 520(7546):186–190.

36. Tsai SQ, et al. (2015) GUIDE-seq enables genome-wide profiling of off-target cleavage by CRISPR-Cas nucleases. Nat Biotechnol 33(2):187–197.

37. Lin Y, et al. (2014) CRISPR/Cas9 systems have off-target activity with insertions or deletions between target DNA and guide RNA sequences. Nucleic Acids Res 42(11):7473–7485.

38. Semenova E, et al. (2011) Interference by clustered regularly interspaced short palindromic repeat (CRISPR) RNA is governed by a seed sequence. Proc Natl Acad Sci U S A 108(25):10098–10103.

39. Dong D, et al. (2016) The crystal structure of Cpf1 in complex with CRISPR RNA. Nature:1–16.

40. Hayes RP, et al. (2016) Structural basis for promiscuous PAM recognition in type I-E Cascade from E. coli. Nature 530(7591):499–503.

41. Jiang F, et al. (2016) Structures of a CRISPR-Cas9 R-loop complex primed for DNA cleavage. Science (80-) 8282(January):1–8.

42. Jiang F, Zhou K, Ma L, Gressel S, Doudna JA (2015) A Cas9-guide RNA complex preorganized for target DNA recognition. Science (80-) 348(6242):1477–1481.

43. Jinek M, et al. (2014) Structures of Cas9 endonucleases reveal RNA-mediated conformational activation. Science 343(6176):1247997.

44. Yamano T, et al. (2016) Crystal Structure of Cpf1 in Complex with Guide RNA and Target DNA. Cell 165(4):949–962.

45. Nishimasu H, et al. (2014) Crystal structure of Cas9 in complex with guide RNA and target DNA. Cell 156(5):935–949.

46. Blosser TR, et al. (2015) Two distinct DNA binding modes guide dual roles of a CRISPR-cas protein complex. Mol Cell 58(1):60–70.

47. Szczelkun MD, et al. (2014) Direct observation of R-loop formation by single RNA-guided Cas9 and Cascade effector complexes. Proc Natl Acad Sci U S A 111(27):9798–803.

48. Rutkauskas M, et al. (2015) Directional R-loop formation by the CRISPR-cas surveillance complex cascade provides efficient off-target site rejection. Cell Rep 10(9):1534–1543.

49. Singh D, Sternberg SH, Fei J, Ha T, Doudna JA (2016) Real-time observation of DNA recognition and rejection by the RNA-guided endonuclease Cas9. Nat Commun 7:48371.

50. Sternberg SH, Redding S, Jinek M, Greene EC, Doudna JA (2014) DNA interrogation by the CRISPR RNA-guided endonuclease Cas9. Nature 507(7490):62–67.

51. Josephs EA, et al. (2015) Structure and specificity of the RNA-guided endonuclease Cas9 during DNA interrogation, target binding and cleavage. Nucleic Acids Res 43(18):8924–41.

52. Jo MH, et al. (2015) Human Argonaute 2 Has Diverse Reaction Pathways on Target RNAs. Mol Cell 59(1):117–124.

53. Salomon WE, Jolly SM, Moore MJ, Zamore PD, Serebrov V (2015) Single-Molecule Imaging Reveals that Argonaute Reshapes the Binding Properties of Its Nucleic Acid Guides. Cell 162(1):84–95.

54. Slaymaker IM, et al. (2015) Rationally engineered Cas9 nucleases with improved specificity. Science (80-) 351(6268):84–88.

55. Kleinstiver BP, et al. (2015) Engineered CRISPR-Cas9 nucleases with altered PAM specificities. Nature 523(7561):481–485.

56. Kleinstiver BP, et al. (2015) Broadening the targeting range of Staphylococcus aureus CRISPR-Cas9 by modifying PAM recognition. Nat Biotechnol 33(November):1–7.

57. Kleinstiver BP, et al. (2016) High-fidelity CRISPR-Cas9 nucleases with no detectable genome-wide off-target effects. Nature 529(7587):490–495.

58. Künne T, Swarts DC, Brouns SJJ (2014) Planting the seed: Target recognition of short guide RNAs. Trends Microbiol 22(2):74–83.

59. Boyle EA, et al. (2016) High-throughput biochemical profiling reveals Cas9 off-target binding and unbinding heterogeneity. bioRxiv.

60. Fu Y, Sander JD, Reyon D, Cascio VM, Joung JK (2014) Improving CRISPR-Cas nuclease specificity using truncated guide RNAs. Nat Biotechnol 32(3):279–284.

61. Ran FA, et al. (2013) Double nicking by RNA-guided CRISPR cas9 for enhanced genome editing specificity. Cell 154(6):1380–1389.

62. Haeussler M, et al. (2016) Evaluation of off-target and on-target scoring algorithms and integration into the guide RNA selection tool CRISPOR. Genome Biol 17(1):148.

63. Bae S, Park J, Kim J (2014) Sequence analysis Cas-OFFinder: a fast and versatile algorithm that searches for potential off-target sites of Cas9 RNA-guided endonucleases. Bioinformatics 30(10):1473–1475.

64. Labun K, Montague TG, Gagnon JA, Thyme SB, Valen E (2016) CHOPCHOP v2: a web tool for the next generation of CRISPR genome engineering. Nucleic Acids Res 44(May):W272–W276.

65. Heigwer F, Kerr G, Boutros M (2014) E-CRISP: fast CRISPR target site identification. Nat Methods 11(2):122–123.

66. MIT CRISPR design tool Available at: http://crispr.mit.edu:8079/about.

67. Listgarten J, Weinstein M, Elibol M, Hoang L, Doench J (2016) Predicting off-target effects for end-to-end CRISPR guide design.bioRx/v:1–16.

68. Agarwal V, Bell GW, Nam JW, Bartel DP (2015) Predicting effective microRNA target sites in mammalian mRNAs. Elife 4(AUGUST2015):1–38.

69. Farasat I, Salis HM (2016) A Biophysical Model of CRISPR/Cas9 Activity for Rational Design of Genome Editing and Gene Regulation. PLoS Comput Biol 12(1):1–33.

70. Khorshid M, Hausser J, Zavolan M, van Nimwegen E (2013) A Biophysical miRNA-mRNA Interaction Model Infers Canonical and Noncanonical Targets. Nat Methods 10(3):253–255.

71. Bisaria N, Jarmoskaite I, Herschlag D (2017) Lessons from Enzyme Kinetics Reveal Specificity Principles for RNA-Guided Nucleases in RNA Interference and CRISPR-Based Genome Editing. Cell Syst 4(1):21–29.

72. Sternberg SH, LaFrance B, Kaplan M, Doudna JA (2015) Conformational control of DNA target cleavage by CRISPR-Cas9. Nature 527(7576):1–14.

73. Leenay RT, et al. (2016) Identifying and Visualizing Functional PAM Diversity across CRISPR-Cas Systems. Mol Cell 62(1):137–147.

74. Xue C, et al. (2015) CRISPR interference and priming varies with individual spacer sequences. Nucleic Acids Res 43(22):10831–10847.

75. Sugimoto N, Nakano SI, Yoneyama M, Honda KI (1996) Improved thermodynamic parameters and helix initiation factor to predict stability of DNA duplexes. Nucleic Acids Res 24(22):4501–4505.

76. Watkins NE, et al. (2011) Thermodynamic contributions of single internal rAdA, rCdC, rGdG and rUdT mismatches in RNA/DNA duplexes. Nucleic Acids Res 39(5):1894–1902.

77. Brouns SJJ, et al. (2008) Small CRISPR RNAs guide antiviral defense in prokaryotes. Science 321(5891):960–4.

78. Xue C, et al. (2016) Conformational Control of Cascade Interference and Priming Activities in CRISPR Immunity Short Article Conformational Control of Cascade Interference and Priming Activities in CRISPR Immunity. Mol Cell:1–9.

79. Marraffini LA, Sontheimer EJ (2010) Self versus non-self discrimination during CRISPR RNA-directed immunity. Nature 463(7280):568–71.

80. Klein M, Chandradoss SD, Depken M, Joo C (2017) Why Argonaute is needed to make microRNA target search fast and reliable. Semin Cell Dev Biol. doi:10.1016/j.semcdb.2016.05.017.

81. Shmakov S, et al. (2015) Discovery and Functional Characterization of Diverse Class 2 CRISPR-Cas Systems. Mol Cell 60(3):385–397.

82. Kaya E, et al. (2016) A bacterial Argonaute with noncanonical guide RNA specificity. Proc Natl Acad Sci U S A 113(15):1524385113-.

83. Novozhilov AS, Georgy PK, Koonin E V (2006) Biological applications of the theory of birth-and-death processes. Brief Bioinform 7(1):70–85.

84. Nowak MA (2006) Evolutionary Dynamics: exploring the equations of life (Harvard University Press).

85. Bartel DP (2009) MicroRNAs: Target Recognition and Regulatory Functions. Cell 136(2):215–233.

86. Swarts DC, et al. (2014) The evolutionary journey of Argonaute proteins. Nat Struct Mol Biol 21(9):743–753.

